# Mitochondrial genomes of leafhoppers and a parasitoid fly expand genomic resources for the Eastern Canadian Arctic

**DOI:** 10.64898/2026.10.13.759010

**Authors:** Florent Sylvestre, Nicolas Plante, Joshua Molligan, Amélie Grégoire Taillefer, Edel-Pérez-López

## Abstract

Arctic and Subarctic insect biodiversity remains poorly represented in molecular reference databases, limiting the identification of resident taxa and the detection of ecological changes associated with climate warming. Leafhoppers are particularly relevant to northern surveillance because they are sensitive to changes in vegetation and climate, and some species transmit phytoplasmas and other plant pathogens. We used Illumina sequencing to characterize mitochondrial genomes from pooled, morphologically identified leafhoppers collected through the Nunavik Sentinels community science program in Eeyou Istchee, Nunavik, and Nunatsiavut. Eight complete mitochondrial genomes were recovered, representing seven leafhopper lineages and one pipunculid fly. The genomes ranged from 15,475 to 18,051 bp, had mean sequencing depths of 1,003×–3,366×, and contained the 37 genes expected in insect mitochondria. Molecular analyses revealed a *Sorhoanus*-like lineage within a pool identified as *Psammotettix alienus*, demonstrating the capacity of untargeted sequencing to detect taxa overlooked during morphological sorting. A *Tomosvaryella*-like mitochondrial genome was also recovered from a pool composed of *Diplocolenus abdominalis*. This finding is consistent with the presence of parasitoid within a host but does not establish a direct host–parasitoid association. COI comparisons supported the higher level placement of the taxa but frequently failed to provide species-level resolution because of incomplete reference coverage, inconsistent nomenclature, and low divergence among closely related species. Phylogenetic analysis of the 13 mitochondrial protein-coding genes supported the expected tribal placements, but provided limited resolution for some deeper relationships, reinforcing the need to expand the availability of nuclear genomic data for leafhopper phylogenetics. These voucher-linked and geographically referenced mitogenomes expand the molecular representation of northern insects and provide foundational resources for monitoring leafhopper diversity, potential plant disease vectors, and their natural enemies in rapidly changing northern ecosystems.

## 1 INTRODUCTION

Biodiversity monitoring increasingly depends on the availability of reliable molecular reference resources. Such resources support species identification, the reconstruction of evolutionary and biogeographic histories and the detection of changes in biological communities (Brandies et al. 2019; Hogg et al. 2022; Hogg et al. 2025). This need is particularly pressing for insects, which account for a substantial proportion of terrestrial biodiversity but are undergoing rapid and spatially heterogeneous changes in abundance, distribution, and community composition (Wagner 2020).

Mitochondrial DNA has played a central role in the development of molecular ecology (Ballard and Whitlock 2004), with population genetic, phylogeography and environmental DNA studies relying on mitochondrial markers. The Cytochrome *c* oxidase subunit I (COI) gene have been largely used to study recent evolutionary history, and has become the standard barcode for animal identification, with reference databases such as the Barcode of Life Data System linking morphological identification to molecular sequences (BOLD; Ratnasingham and Hebert 2007; Adamowicz 2015). The availability of complete mitochondrial reference genomes has only strengthened the importance of the mitochondria for molecular ecology, consolidating population genetic and phylogenetic studies ((Ballard and Whitlock 2004; Xie et al. 2025) and expanding species barcodes identification beyond the COI for species complex or low divergence taxa (Romão et al. 2026).

Although thousands of arthropod barcodes are publicly available, many lineages and geographic regions remain poorly represented. This is especially important for morphologically challenging groups such as insects, for which species-level identification requires highly specialized expertise and the examination of adult male genitalia (Beirne 1956; Godfray 2002). Females, immature stages, damaged specimens, and organisms recovered from bulk or environmental samples may therefore remain unidentified without representative molecular references. Building geographically explicit barcode and mitogenome libraries is a prerequisite for developing scalable molecular insect monitoring (Seidensticker et al. 2026).

These limitations are particularly consequential in Arctic and Subarctic ecosystems. Northern environments are undergoing rapid climatic and ecological transformation, but uncertainty in species occurrence and the scarcity of long-term observations make changes in arthropod communities difficult to quantify (Gillespie et al. 2020; Høye et al. 2021; Grégoire Taillefer et al. 2024). Molecular resources generated from well-identified northern specimens would help overcome this limitation and are essential for establishing the longitudinal surveillance systems needed to evaluate how climate change is reshaping northern insect communities.

Leafhoppers (Hemiptera: Cicadellidae) are particularly relevant to northern biodiversity surveillance. With more than 20,000 described species worldwide and over 1,200 reported from Canada and Alaska, Cicadellidae is among the largest families of herbivorous insects (Weintraub and Beanland 2006). Their distributions are closely associated with those of their host plants, making them potentially sensitive indicators of changes in vegetation and climate. Several species are economically important pests or vectors of phytoplasmas, viruses, and bacteria that cause plant diseases (Olivier et al. 2014; Brochu et al. 2023; Santos et al. 2024). Changes in their abundance, phenology, and geographic distribution could therefore affect not only insect biodiversity but also plant health, agriculture, and northern food systems. Some pest and vector species are predicted to expand northward under warmer climatic scenarios, underscoring the need for molecular reference resources capable of distinguishing resident Arctic taxa from newly arriving species (Santos et al. 2024; Santos et al. 2026).

Building on our collaboration with the Nunavik Sentinels program, we examined leafhoppers collected between 2018 and 2026 across Eeyou Istchee and Nunavik in Québec and Nunatsiavut in Labrador. Although the program primarily monitors pollinators, its collections contain a broad diversity of nontarget insects from Boreal, Subarctic, and Arctic environments. Previous examination of this material revealed an underdocumented assemblage of leafhoppers and established the first baseline for their diversity in the Eastern Canadian Arctic (Santos et al. 2026). These geographically referenced specimens also provide an opportunity to develop molecular resources for the long-term monitoring of northern insect communities.

In this study, we used short-read Illumina sequencing to assemble and annotate seven complete mitochondrial genomes from leafhopper species collected in the Eastern Canadian Arctic. Unexpectedly, sequencing also revealed the complete mitochondrial genome of a parasitoid fly of the genus *Tomosvaryella*, likely originating from parasitized leafhoppers. We compared the genomic structure and composition of these mitogenomes with available mitochondrial sequences and evaluated their phylogenetic placement. This resource is foundational for monitoring Arctic leafhopper communities, detecting potential plant disease vectors, and investigating parasitoid–host associations relevant to natural and biological control.

## 2 MATERIALS AND METHODS

### 2.1 Insect collection and genomic sequencing

Insects were collected through the Nunavik Sentinels community science program between 2018 and 2023 in the Eeyou Istchee and Nunavik regions of Québec and between 2022 and 2023 in Nunatsiavut, Newfoundland and Labrador. Detailed descriptions of the sampling program, collection protocols, and taxonomic identification procedures are provided in Grégoire-Taillefer et al. (2024) and Santos et al.(2026). Obtaining enough high-quality DNA from individual specimens was challenging because of their small body size, approximately 1–3 mm for the smallest species, and the partial degradation of DNA during collection and storage. Genomic DNA was therefore extracted from pools of 8 to 10 morphologically identified individuals as previously described (Molligan et al. 2025).

Sufficient numbers of specimens were available for seven target species: *Diplocolenus abdominalis* (Fabricius, 1803), *Hebecephalus algidus* (DeLong & Davidson, 1935), *Macustus grisescens* (Zetterstedt, 1828), *Psammotettix alienus* (Dahlbom, 1850), *Limotettix myralis* (Medler, 1958), *Limotettix plutonius* (Uhler, 1877), and *Stroggylocephalus mixtus* (Say, 1825) (**Figure 1**). At the time of the analysis, no complete mitochondrial reference genome was available in the National Center for Biotechnology Information (NCBI) database for any of these species. Genomic DNA was submitted to the *Centre d’expertise et de services Génome Québec* (Montréal, Québec, Canada) for quality control, library preparation using the NEBNext Ultra II DNA Library Prep Kit, and sequencing on an Illumina NovaSeq X platform. Sequencing targeted approximately 30 million 150-bp paired-end reads per pool of samples.

**Figure 1.**
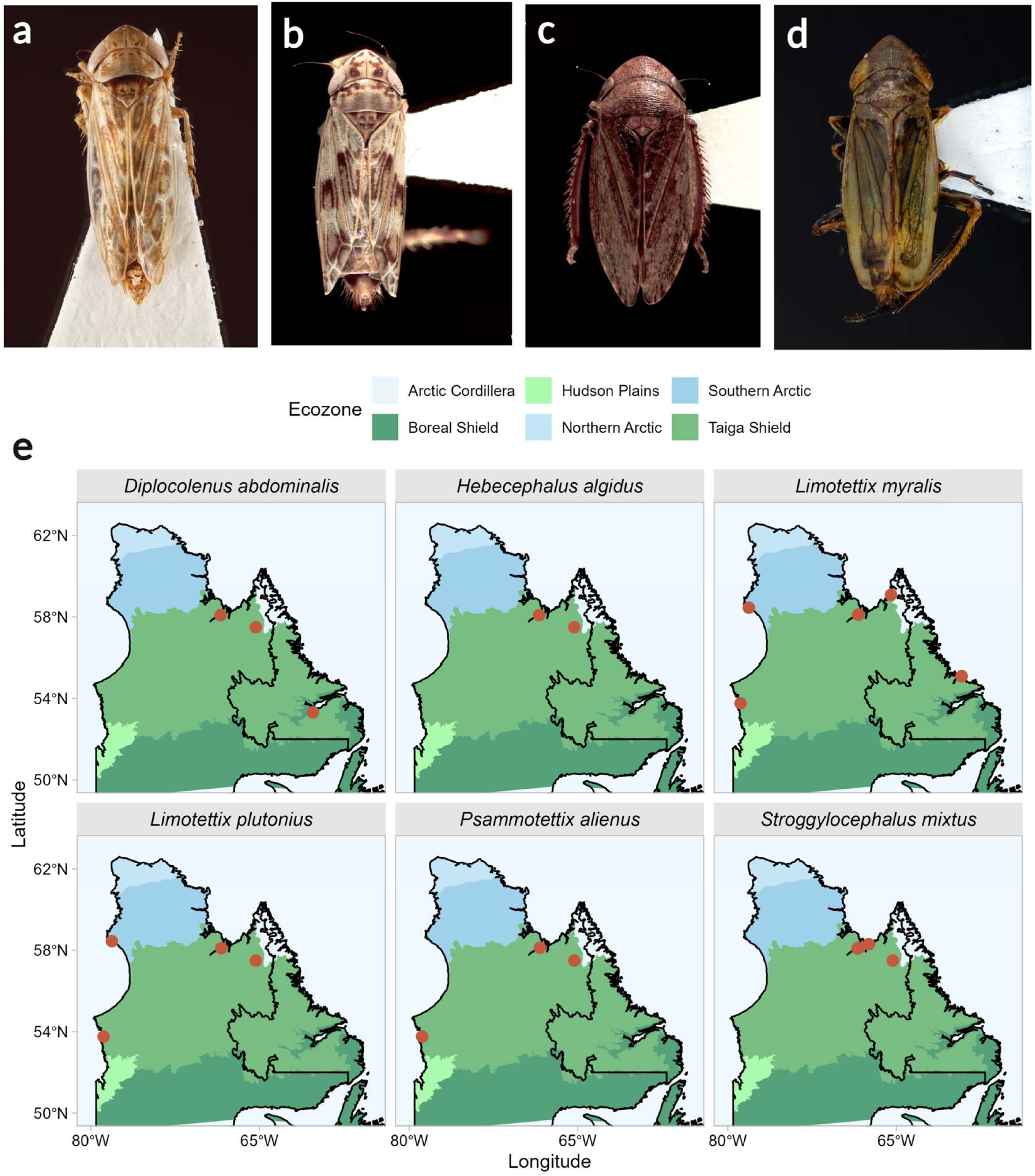
Representative leafhoppers and geographic distribution of the focal species in the Eastern Canadian Arctic. (a–d) Dorsal views of representative specimens of (a) *Psammotettix alienus*, (b) *Hebecephalus algidus*, (c) *Stroggylocephalus mixtus*, and (d) *Diplocolenus abdominalis*. (e) Collection localities of *Diplocolenus abdominalis*, *Hebecephalus algidus*, *Limotettix myralis*, *Limotettix plutonis*, *Psammotettix alienus*, and *Stroggylocephalus mixtus*. Orange circles indicate collection sites for each species previously reported by Santos-Almeida et al. (2026). Background colours represent the six ecozones shown in the legend: Arctic Cordillera, Boreal Shield, Hudson Plains, Northern Arctic, Southern Arctic, and Taiga Shield.

### 2.2 Mitochondrial genome assembly

Raw sequencing reads were processed using fastp v0.15.0 (Chen et al. 2018) to remove Illumina adapters and low-quality reads. Reads were discarded when more than 40% of their bases had Phred quality scores below 15. Sequence quality was evaluated before and after filtering using FastQC (https://www.bioinformatics.babraham.ac.uk/projects/fastqc/).

Mitochondrial reads were identified and assembled using the get_organelle_from_reads.py workflow implemented in GetOrganelle v1.7.7.1 (Jin et al. 2020). Assemblies were generated using the animal_mt reference database, 10 extension rounds, and k-mer sizes of 21, 45, 65, 85, and 105. Because DNA was extracted from pools of multiple individuals, intraspecific mitochondrial variation frequently produced complex assembly graphs which prevented the automatic recovery of a single contiguous mitochondrial sequence. Assembly graphs were consequently inspected and curated in Bandage v0.8.1 (Wick et al. 2015). For each sample, the path supported by the highest read coverage was retained to generate a representative consensus sequence. Alternative paths consistent with lower-frequency haplotypes and unsupported dead ends produced by incomplete reconstruction were removed. Curated graphs were subsequently processed using the get_organelle_from_assembly.py script to recover the final mitochondrial genome sequences.

### 2.3 Genome annotation and taxonomic validation

Consensus mitochondrial genomes were annotated using MITOS2 v2.1.10+galaxy0 (Bernt et al. 2013) and the RefSeq 89 Metazoa reference dataset. Protein-coding genes, transfer RNA, and ribosomal RNA were predicted using the invertebrate mitochondrial genetic code. Circular genome sequences were standardized by relocating the sequence origin immediately upstream of *trnI*, ensuring that the *cox1* gene was located on the forward strand.

Transfer RNA predictions were complemented using tRNAscan-SE v2.0 (Chan et al. 2021), with a minimum score of 5. Predictions located within 5 bp of the corresponding MITOS2 annotation were considered concordant and merged, and tRNA annotations with scores below 15 were removed. Protein-coding gene boundaries were further evaluated using custom Python scripts and Geneious Prime v2025.1.3 (https://www.geneious.com). Alignments with homologous mitochondrial genes from related taxa were examined to refine start and stop codons and resolve inconsistencies between automated annotations.

Taxonomical identifications were assessed using the Barcode of Life Data System identification engine (BOLD) (Ratnasingham et al. 2024). The COI sequence recovered from each mitochondrial genome was queried against the BOLD Public Record Barcode Database using the Species Level Barcode Records identification mode (Ratnasingham and Hebert 2013). Because some focal species were not represented by validated COI reference sequences, BOLD results were interpreted primarily as independent support for genus- or tribe-level placement instead of species-level identifications.

### 2.4 Phylogenetic validation

To evaluate the phylogenetic placement of the newly assembled mitochondrial genomes, we retrieved the sequences and annotations of 39 publicly available Deltocephalinae mitochondrial genomes from NCBI. Reference taxa were selected based on previous phylogenetic analyses of the subfamily (**Table S1**) (Zahniser and Dietrich 2008; Molligan et al. 2025).

The amino acid sequences of the 13 mitochondrial protein-coding genes were extracted from the downloaded references and newly assembled genomes using a custom Python script. Sequences for each protein were aligned independently in MAFFT v7.525 using the L-INS-i algorithm and 1,000 refinement iterations (Katoh and Toh 2008). Poorly aligned positions were removed using Gblocks v1.0 with default parameters (Castresana 2000). The trimmed alignments were then concatenated into a single amino acid matrix. Maximum-likelihood phylogenetic inference was performed using IQ-TREE v2.1.3 (Minh et al. 2020). ModelFinder Plus was used to identify the best-fitting substitution model, and branch support was assessed with 1,000 ultrafast bootstrap replicates. The resulting phylogeny was visualized and annotated using Interactive Tree of Life v6 (iTOL) (Letunic and Bork 2024).

### 2.5 Read coverage and mitochondrial variation

Quality-filtered reads were mapped back to the corresponding mitochondrial genome assemblies using BWA-MEM v0.7.18 (Li 2013) with default parameters. For samples containing more than one mitochondrial genome, reads were mapped against a combined reference containing all recovered mitochondrial sequences. This approach reduced erroneous cross-mapping between related sequences present in the same pooled sample. Sequencing depth was summarized in 500-bp windows using mosdepth v0.3.12 (Pedersen and Quinlan 2018). Polymorphism were identified from the alignments using mpileup in bcftools v1.21 (Li 2011), filtering out variations supported by less than 10% of the total coverage. Circular genome maps integrating gene annotations, read coverage, variant density, and GC content in 500-bp windows were generated using Proksee (Grant et al. 2023). Nucleotide-level synteny among mitochondrial genomes based on BLAST comparisons was visualized using pyGenomeViz (Shimoyama 2024). All remaining statistical analyses and visualizations were performed in R (Wickham et al. 2019; R Core Team 2021).

## 3 RESULTS AND DISCUSSION

### 3.1 Recovery of eight complete mitochondrial genomes from Arctic insect samples

Complete mitochondrial genome sequences were recovered for six of the seven target leafhopper species. Assembly of the *M. grisescens* mitochondrial genome was unsuccessful because GetOrganelle identified only 2,896 mitochondrial reads from this library (**Table S2**). The remaining target-species assemblies had sufficient read depth to recover complete circular mitochondrial genomes, although six required manual resolution of the genome graph.

In addition to the six expected leafhopper mitochondrial genomes, two unexpected genomes were recovered from the pooled samples: (*i*) a second leafhopper mitochondrial genome closely related to the genus *Sorhoanus* and (*ii*) a mitochondrial genome from a pipunculid fly. The final dataset therefore comprised eight complete mitochondrial genomes, seven from leafhoppers and one from a parasitoid fly.

The assembled genomes ranged from 15,475 to 18,051 bp, with mean sequencing depths ranging from 1,003× to 3,366× (**Table 1**). All eight genomes contained the 37 genes expected in a typical insect mitochondrial genome: 13 protein-coding genes, 22 tRNA genes, and two rRNA genes (**Table 1**, **Figures 2**, **Figure 3**). Their completeness, conserved gene content, and high sequencing depth support the overall quality of the assemblies despite the additional graph-curation steps required for pooled samples.

**Figure 2.**
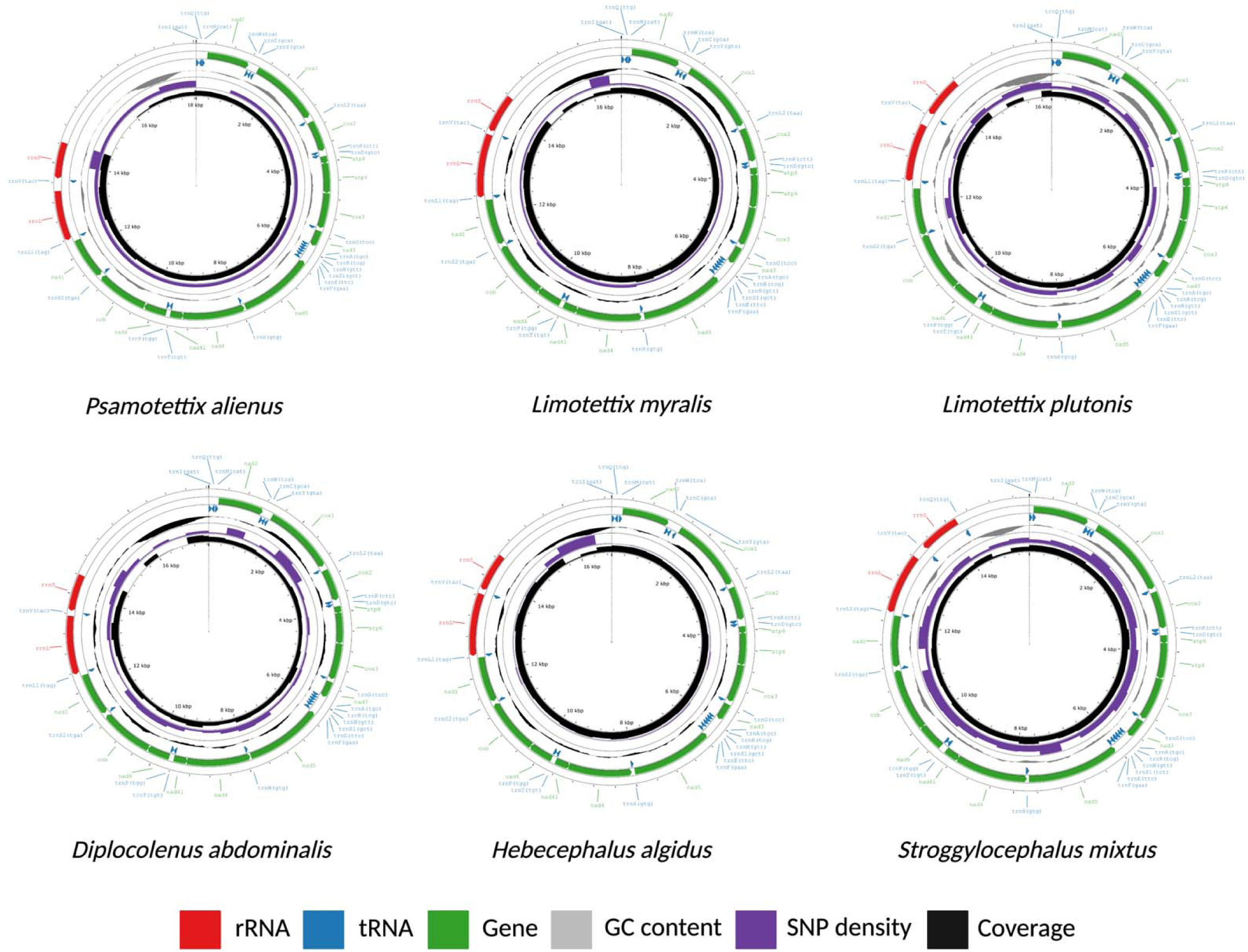
Mitochondrial genome organization of six leafhopper species collected in northern Québec. Circular maps show the mitochondrial genomes of *Psammotettix alienus*, *Limotettix myralis*, *Limotettix plutonis*, *Diplocolenus abdominalis*, *Hebecephalus algidus*, and *Stroggylocephalus mixtus*. Protein-coding genes, transfer RNA genes, and ribosomal RNA genes are shown in green, blue, and red, respectively, with arrows indicating transcriptional orientation. Inner tracks show GC content (grey), single-nucleotide polymorphism (SNP) density (purple), and sequencing coverage (black). Genome coordinates are indicated in kilobase pairs.

**Figure 3.**
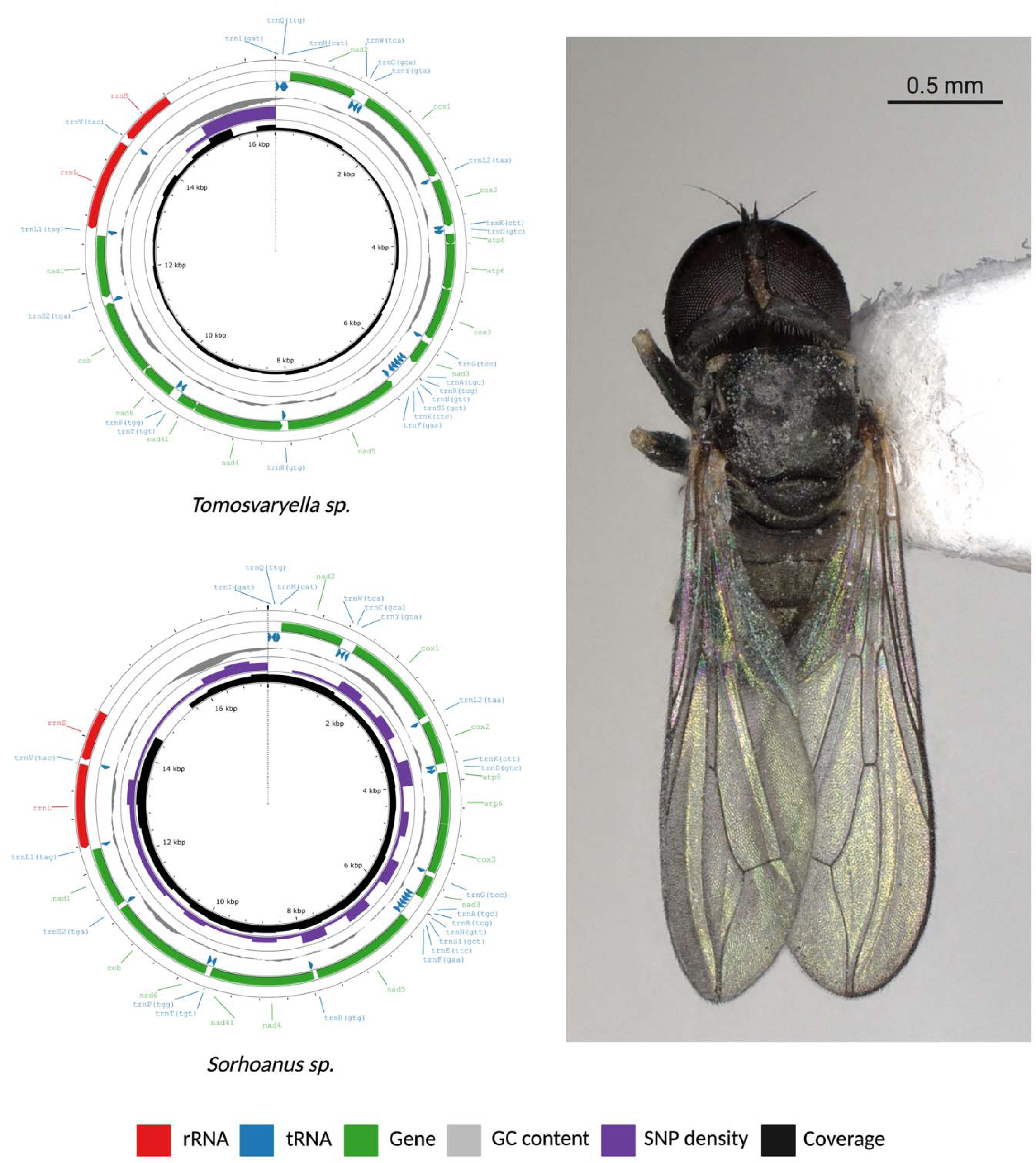
Mitochondrial genomes of Tomosvaryella sp. and Sorhoanus sp. recovered in this study. Circular maps show the mitochondrial genomes of *Tomosvaryella* sp. (upper left) and *Sorhoanus* sp. (lower left). Protein-coding genes, transfer RNA genes, and ribosomal RNA genes are shown in green, blue, and red, respectively, with arrows indicating transcriptional orientation. Inner tracks represent GC content (grey), single-nucleotide polymorphism (SNP) density (purple), and sequencing coverage (black). Genome coordinates are indicated in kilobase pairs. The photograph on the right shows a dorsal view of *Tomosvaryella* sp. reared in our lab. Scale bar, 0.5 mm.

**Table 1.**
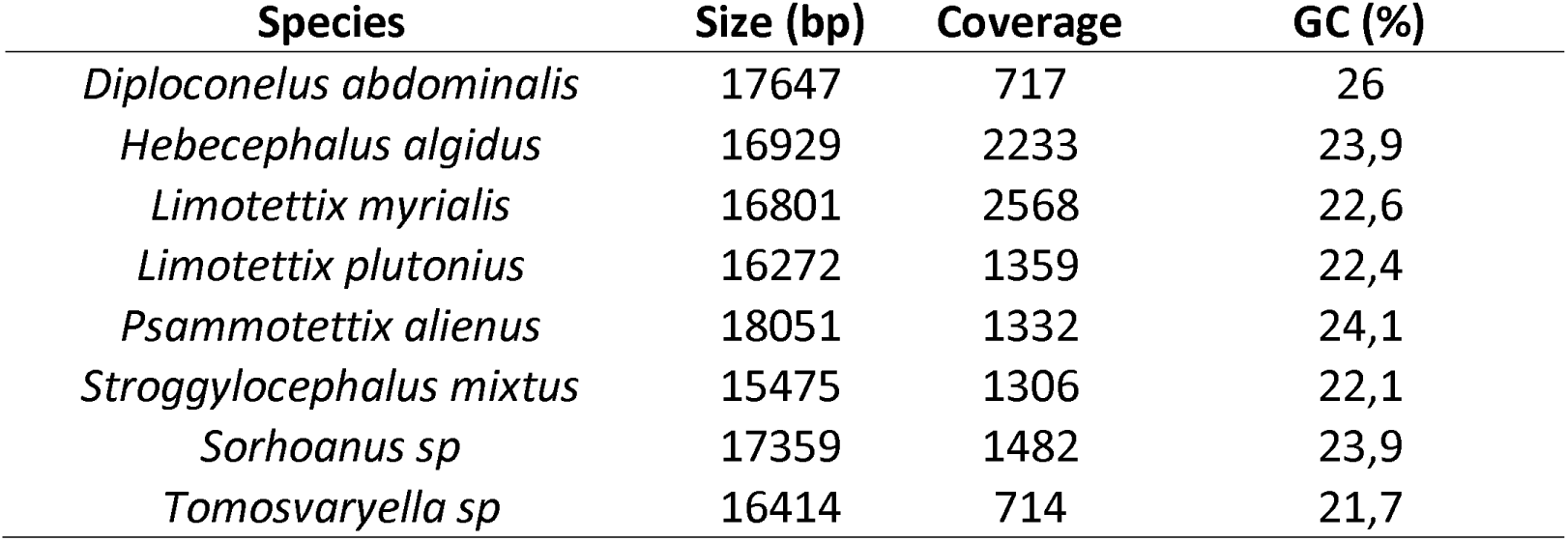
Main statistics of assembled mitogenomes in this study.

### 3.2 Molecular data revealed an unexpected leafhopper lineage

Morphological identification of leafhoppers is often difficult because many species share similar external characteristics and reliable identification may require examination of male genitalia. Although the specimens used here were examined by researchers with experience in leafhopper taxonomy, molecular analysis of the *P. alienus* pool unexpectedly recovered a second complete leafhopper mitochondrial genome. Its COI sequence shared more than 98% identity with sequences assigned to the genus *Sorhoanus* in BOLD (**Figure 3A**).

This finding illustrates both the value and the limitations of molecular identification. The genus *Sorhoanus* was not reported in our recent baseline survey of leafhoppers from the eastern Canadian Arctic, in which we identified 21 genera (Santos et al. 2026). Its detection here resulting from a mixed-species sample suggests that *Sorhoanus* specimens were likely present at low abundance and accidently pooled with morphologically similar specimens during sorting. Sequence-based methods can reveal taxonomic diversity that may remain undetected when rare specimens are overlooked or mixed with morphologically similar taxa. However, the interpretation of molecular data depends on the accuracy and completeness of the reference databases. Molecular approaches should therefore complement and not replace taxonomic expertise. Vouchered specimens, expert identifications, and well-curated reference sequences remain indispensable for resolving difficult insect groups and improving the reliability of barcode databases (Godfray 2002; Santos and Pérez-López 2026).

### 3.3 Detection of a potential leafhopper parasitoid

A complete mitochondrial genome from a pipunculid fly was also recovered unexpectedly from one of the pooled leafhopper samples, belonging to *D. abdominalis* (**Figure 3B**). Its COI sequence showed greater than 90% identity to sequences assigned to the genus *Tomosvaryella* in BOLD. Pipunculidae, commonly known as big-headed flies, include endoparasitoids of Auchenorrhyncha, and members of this family are among the principal parasitoids reported from leafhopper communities (Waloff and Jervis 1987; Kirk-Spriggs and Sinclair 2017).

Although *Tomosvaryella* is one of the most species-rich genera of Pipunculidae, its host associations and biology remain remarkably poorly documented. The few available records of well characterized biocontrol include *Tomosvaryella lepidipes* (Hardy, 1943) developing in adults of the geminate leafhopper *Colladonus geminatus* (Van Duzee, 1951) and *Tomosvaryella vagabunda* (Knab, 1915) attacking the beet leafhopper *Circulifer tenellus* (Baker, 1896) in western North America (Kaloostian 1953; Clarke et al. 2024). More recently, *Tomosvaryella* DNA was detected in specimens associated with *Amblysellus* leafhoppers collected in Idaho, but these associations were not confirmed through rearing (Strausbaugh et al. 2026). This limited biological information likely reflects the difficulty of detecting parasitized hosts and maintaining pipunculids in captivity. Parasitism often produces no obvious external structure, while larval emergence, soil pupation, diapause, and low adult emergence complicate experimental rearing.

Because the mitochondrial genome was recovered from a pool composed exclusively of specimens identified as *D. abdominalis*, the most biologically informative explanation is that one leafhopper contained developing parasitoid tissue. If confirmed, this would represent a new reproductive host record for *Tomosvaryella* and, to our knowledge, the first evidence that *D. abdominalis* can support the development of a pipunculid parasitoid. Nevertheless, accidental inclusion of a small adult fly during pooling cannot be excluded, and the molecular data alone cannot establish a direct association. Confirming this association will require recovering *Tomosvaryella* directly from individually identified *D. abdominalis* specimens. The complete mitochondrial genome reported here provides a reference for these surveys and illustrates how untargeted sequencing can reveal poorly documented natural enemies of northern leafhopper communities, potentially supporting future assessments of their contribution to natural regulation and biological control.

### 3.4 Taxonomic resolution of COI barcoding varied among leafhopper lineages

Comparison of the recovered COI sequences with BOLD records supported the genus-level identification of *D. abdominalis*, *S. mixtus*, *H. algidus*, and *P. alienus* (**Table S3**). However, species-level confirmation was not possible because validated reference sequences were not available. The COI sequence assigned morphologically to *S. mixtus* was identical to a BOLD sequence identified as *Stroggylocephalus placidus* (Provancher, 1889), another species reported from Canada. This result could indicate low mitochondrial divergence between the two species or an inaccurate identification in either the present dataset or the database. However, we tried to trace the precedence of that other material, and it was impossible although it seems to come from a museum (deWaard et al. 2019). Additional analyses will be required to distinguish among these possibilities.

Taxonomic resolution was particularly limited within the tribe Limotettigini. Although the recovered sequences were consistently assigned to this tribe, they showed greater than 97% identity to reference records attributed to the genera *Limotettix*, *Scleroracus*, and *Ophiola*. One sequence was identical to a BOLD record identified as *Ophiola corniculus* (Marshall, 1866), whereas another matched *Scleroracus arctostaphyli* (Ball, 1899) with 100% identity (**Tables S3** and **Table S4**). These apparently conflicting assignments likely reflect differences in the taxonomic frameworks used to identify and deposit the reference material. The genus *Ophiola*, for example, was synonymized with *Scleroracus* by Oman (1949), whereas *Scleroracus* has subsequently been treated either as a distinct genus or as a subgenus of *Limotettix*, with *S. arctostaphyli typically known as Limotettix arctostaphyli* (Ball, 1899) (Ribaut 1986; Hamilton 1994; McKamey 2001; Tishechkin 2019). Consequently, closely related or identical reference sequences may be deposited under different generic combinations, limiting genus- and species-level identification based solely on sequence similarity.

These results illustrate how taxonomic errors and incomplete representation can propagate uncertainty within barcode databases (Virgilio et al. 2010; Fujisawa and Imai 2026). They also reflect intrinsic limitations of mitochondrial barcoding. Although COI successfully discriminates many leafhopper species, its resolution can be reduced by mitochondrial introgression, incomplete lineage sorting, low evolutionary rates, or recent divergence (Virgilio et al. 2010; Watanabe et al. 2024). Previous studies have successfully applied COI barcoding to leafhopper identification ((Kamitani 2011; Foottit et al. 2014; Zhang et al. 2019; Emam et al. 2020), but approximately 25% of Nearctic Auchenorrhyncha species examined by Foottit et al. (2014) showed less than 1% interspecific COI divergence. Reliable molecular monitoring of Arctic leafhoppers will therefore require continued sampling and the development of voucher-linked regional reference libraries.

### 3.5 Mitochondrial genome organization and evolution

Gene content and overall genome organization were highly conserved among the newly assembled leafhopper mitochondrial genomes and were generally consistent with previously characterized Cicadellidae mitogenomes (Xie et al. 2024; Molligan et al. 2025). The mitochondrial genome of the pipunculid fly also displayed the gene complement expected for insects. In all assemblies, the region corresponding to the putative mitochondrial control region had comparatively low GC content (**Table 1**). Differences in the length of this noncoding region accounted for much of the variation in total genome size among taxa (**Figures 2** to **Figure 4**). The principal difference in gene organization involved *trnQ* in *S. mixtus*. In this genome, *trnQ* was located on the reverse strand near the small ribosomal RNA gene (*rrnS*), instead of in its typical position on the forward strand between *trnI* and *trnM* (**Figures 2** to **Figure 4**). This arrangement may represent a lineage-specific tRNA translocation, although validation with additional specimens or long-read sequencing would help exclude an assembly artifact. Consistent with the invertebrate mitochondrial genetic code, the most frequently observed initiation codons were ATT and ATG, while ATC, ATA, and TTG occurred less frequently. Complete TAA and TAG termination codons and truncated T stop codons were observed among the protein-coding genes (**Table 2**).

**Figure 4.**
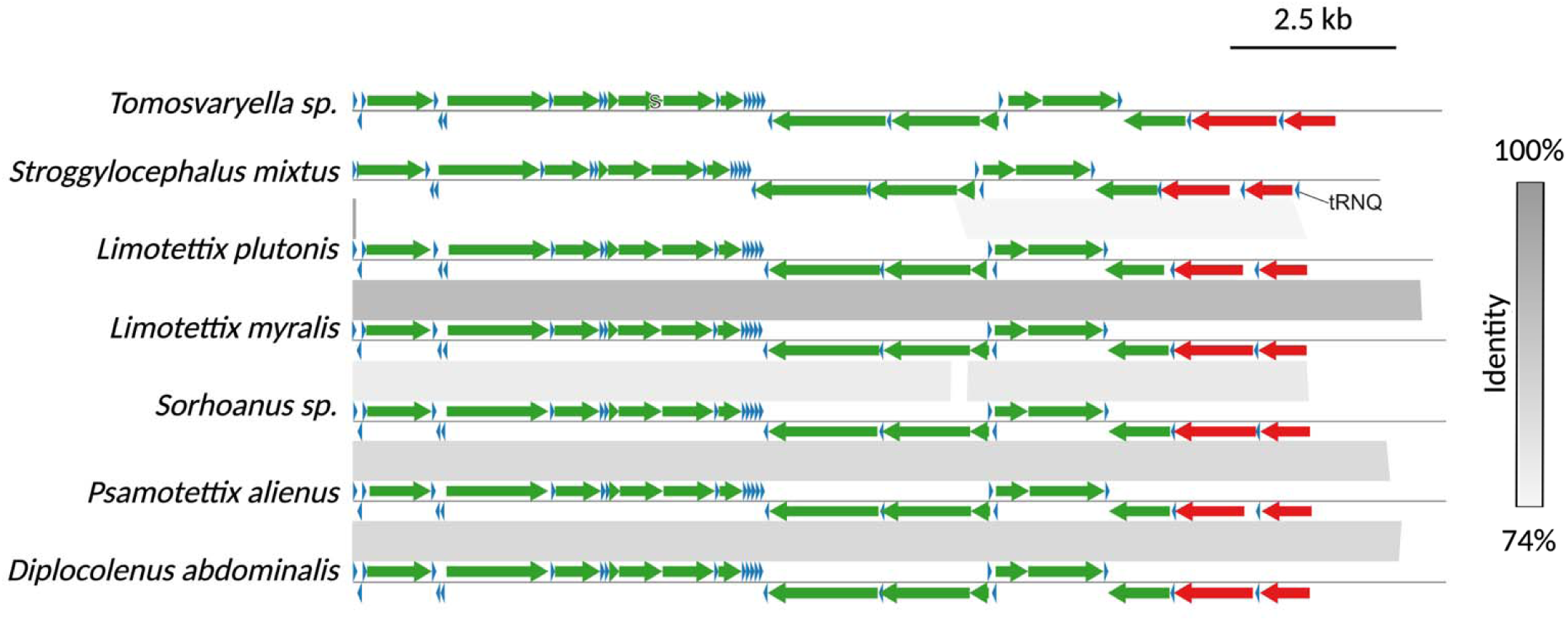
Comparison of mitochondrial gene organization and sequence similarity among seven arthropod taxa. Linearized mitochondrial genomes are shown for *Tomosvaryella* sp., *Stroggylocephalus mixtus*, *Limotettix plutonis*, *Limotettix myralis*, *Sorhoanus* sp., *Psammotettix alienus*, and *Diplocolenus abdominalis*. Arrows represent protein-coding genes (green), transfer RNA genes (blue), and ribosomal RNA genes (red); arrow direction indicates transcriptional orientation. Grey ribbons connect homologous regions between adjacent genomes, with shading indicating nucleotide sequence identity from 74% to 100%. The *trnQ* gene is labelled in *S. mixtus*. Scale bar, 2.5 kb.

**Table 2.**
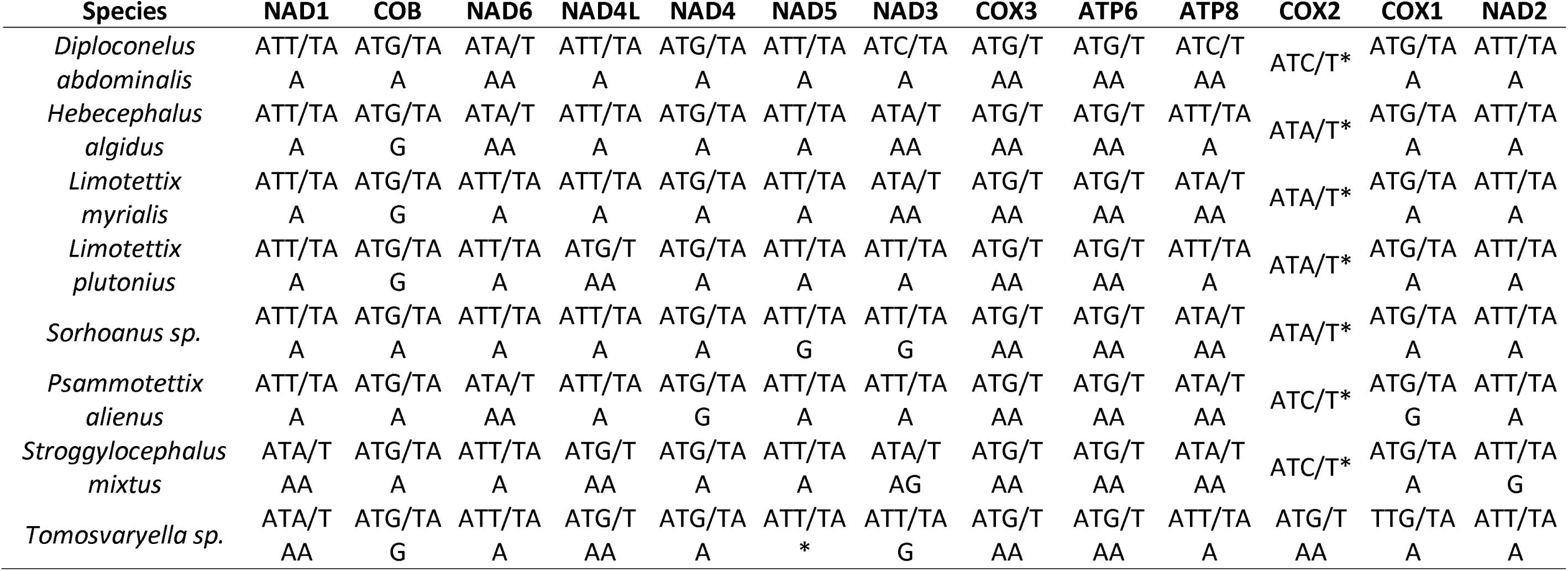
Start and Stop codon usage in mitochondrial protein coding genes.

### 3.6 Phylogenetic placement of the newly sequenced leafhoppers

Phylogenetic reconstruction based on the concatenated amino acid sequences of the 13 mitochondrial protein-coding genes selected mtZOA+F+R6 as the best-fitting evolutionary model according to both the Akaike and Bayesian information criteria. The resulting topology was consistent with the mitochondrial phylogeny reported by Xie et al. (2025), and place the newly assembled leafhopper mitogenomes within the tribes expected from their morphological identifications, supporting their higher-level taxonomical identification.

The sampled tribes were generally recovered as monophyletic. As reported by Xie et al. (2025), *Orosius orientalis* clustered with Cicadulini rather than Opsiini. In contrast to their analysis, *Graminella nigrinota* properly clustered with Deltocephalini instead of Paralimnini, although with moderate bootstrap support (**Figure 5**). Relationships among several tribes were weakly supported, indicating that mitochondrial protein-coding genes alone provide limited resolution for some deeper relationships within Deltocephalinae. Membracidae was placed within the broader cicadellid lineage as the sister group of Eurymelinae, consistent with recent phylogenomic analyses based on nuclear loci (Dietrich et al. 2017). Collectively, these results support the accuracy of the newly assembled genomes while reinforcing the need for nuclear genomic data to resolve deeper relationships among leafhopper tribes, what has been previously discussed (Xie et al. 2025; Wang et al. 2026).

**Figure 5.**
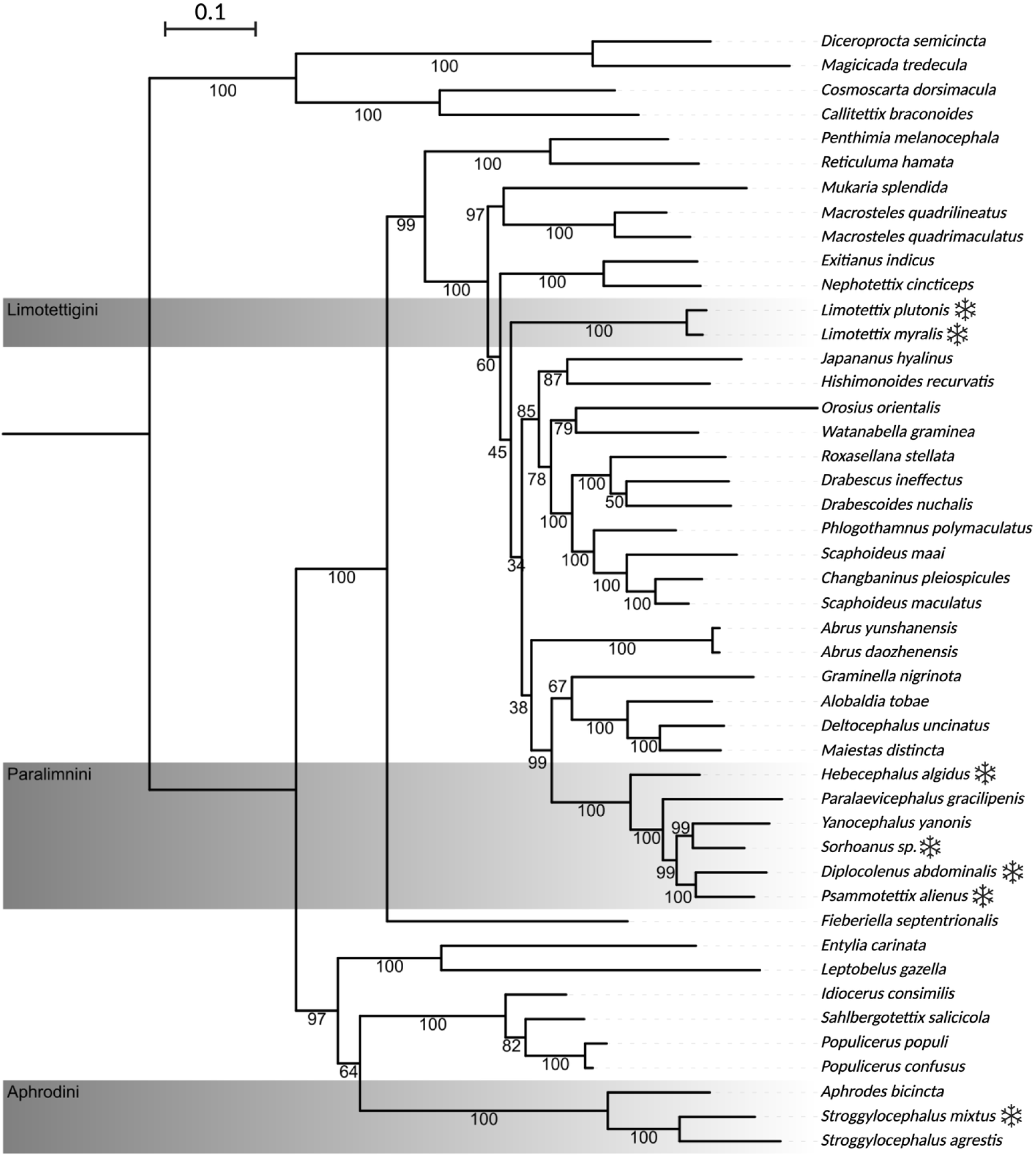
Phylogenetic placement of the leafhopper mitochondrial genomes generated in this study. The phylogeny was inferred using [dataset, inference method, and substitution model]. Snowflake symbols identify taxa whose mitochondrial genomes were generated in this study. Grey bands highlight the tribes Limotettigini, Paralimnini, and Aphrodini. Bootstrap support values indicated. The tree includes *Diceroprocta semicincta*, *Magicicada tredecula*, *Cosmoscarta dorsimacula*, and *Callitettix braconoides* as outgroup. Branch lengths represent estimated substitutions per site; the scale bar corresponds to 0.1 substitutions per site.

## 4 CONCLUSIONS

Nontarget leafhoppers collected through the Nunavik Sentinels program enabled the recovery of eight complete, high-coverage mitochondrial genomes representing seven leafhopper lineages and one pipunculid fly. These genomes address important gaps in the molecular representation of northern insects and provide new references for taxonomic identification, phylogenetic research, and biodiversity monitoring in rapidly changing Arctic and Subarctic environments. This study also exposes the present limitations of molecular identification. Several species lacked validated barcode references, COI did not reliably distinguish some closely related taxa, and an unexpected *Sorhoanus*-like lineage was detected within a morphologically identified sample. Expanding regional reference databases through collaborations between community-based monitoring programs, molecular biologists, and expert taxonomists will therefore be essential for the implementation of reliable, large-scale surveillance.

Leafhoppers are important vectors of phytoplasmas and other plant pathogens, and changes in their distribution could introduce new transmission pathways into northern ecosystems (Plante et al. 2024; Santos et al. 2024). No phytoplasmas were detected in the samples examined here, while studying their microbiome (Pellegrinetti et al. 2026), and there is currently no evidence of leafhopper-transmitted disease affecting cloudberry (*Rubus chamaemorus*) in the study region. Nevertheless, cloudberry and other culturally, ecologically, and economically important northern plants could benefit from proactive surveillance of potential vectors and their associated microorganisms. The mitochondrial references generated here provide a foundation for identifying resident and newly arriving leafhopper species and for evaluating whether environmental change is altering the risk of phytoplasma emergence in northern plant communities.

Finally, the recovery of a *Tomosvaryella* mitochondrial genome demonstrates the potential of untargeted sequencing to detect natural enemies alongside their possible hosts. Although the pooled data do not establish a specific host–parasitoid interaction, the finding is promising because pipunculid flies include specialized parasitoids of leafhoppers and may contribute to the natural regulation of potential disease vectors. This genomic resource provides a foundation for future molecular surveillance of pipunculid parasitoids and their ecological associations. Integrating the monitoring of leafhoppers, their associated microorganisms, and their natural enemies could reveal interactions that influence vector populations and pathogen-transmission risk while informing locally adapted biological control strategies.

## AUTHOR CONTRIBUTIONS

**Florent Sylvestre**: Conceptualization (supporting); writing – original draft (lead), writing – Review and editing (lead) Formal analysis and Data curation (lead); **Nicolas Plante**: – Investigation (support), writing – Review and editing (support); **Joshua Molligan:** Investigation (support), Formal analysis (support), writing – Review and editing (support); **Amélie Grégoire Taillefer**: Investigation (lead), writing – Review and editing (support), **Edel Pérez-López**: Conceptualization (lead), writing – Review and editing (support), funding acquisition (lead).

## FUNDING

The Nunavik Sentinels program, led by the Montréal Insectarium, is made possible through major support from the Royal Bank of Canada (RBC) and the Espace pour la vie Foundation. The Sentinels’ salaries were supported in part by the Canada Summer Jobs and KRG Challenge programs. Air Inuit and Canadian North provided support for air travel. This work was also funded by the Réseau québécois de recherche en agriculture durable (RQRAD), the Ministère de l‘Agriculture, des Pêcheries et de l‘Alimentation du Québec (MAPAQ), and the Fonds de recherche du Québec–Nature et technologies (FRQNT) through the Programme de recherche en partenariat—Agriculture durable—Volet II—2e concours (application number 337847), as well as by the Natural Sciences and Engineering Research Council of Canada (NSERC) through the Alliance-SARI Program (Grant ALLRP 588519-23).

## CONFLICTS OF INTEREST

The authors declare no conflicts of interest associated with this manuscript.

## ACKNOWLEDGEMENTS

We thank Abraão Almeida Santos for his assistance in preparing the maps presented in Figure 1e. The Nunavik Sentinels program thanks Élise Rioux⍰Paquette, formerly of the Kativik Regional Government (KRG), for her commitment and essential contributions to the program’s success. We also thank Patrick Maltais, Isabeau Pratte, and Anna-Kim Pagé-Cornforth at the KRG for their support and expertise. We are grateful to the Arctic Eider Society for integrating the insect component and project updates into the SIKU platform and for their continued collaboration. We extend our thanks to Parcs Nunavik and the communities of Kuujjuaq, Inukjuak, Chisasibi, Happy Valley-Goose Bay, and Makkovik for their participation between 2018 and 2023. We also acknowledge the many collaborators and participants whose involvement made these discoveries possible, including the Sentinels: Toby Nashak, Robert Munick, Tony Anautak, Allison St⍰Cyr, Haylee Petawabano, Destiny Watkin, Robert Andersen, and Olsen Nochasak-Ford. We thank the 2022 organizers at the Nunavik Hunting, Fishing and Trapping Association (Anguvigaq) and participants of the Marralik-Ungunniavik camp; Gregor Gilbert (retired) and Adamie Delisle⍰Alaku at Makivvik Corporation; Erica Oberndorfer, Sean McCann, and Carolyn Parsons at Agriculture and Agri⍰Food Canada; José Gérin⍰Lajoie, Rachel Hussher and Geneviève Dubois at the Université du Québec à Trois⍰Rivières; participants of the Nunami Sukuijainiq-Imaliriijit 2022 marine land camp; youth and organizers of the Pye Centre for Northern Boreal Food Systems camp; as well as the Nunatsiavut Government and the Labrador Inuit Lands. We thank Maxim Larrivée, creator of the Nunavik Sentinels program and Director of the Montréal Insectarium; Julie Jodoin, Director of Espace pour la vie; and Julia Mlynarek, Entomologist at the Insectarium and the Insectarium’s Public Programs and Education Division, for their ongoing support. Also, the Institut de recherche en biologie végétale de Montréal for their administrative support.

## SUPPLEMENTARY FILES

**Table S1.** Complete information of the species included in the phylogenetic analysis and associated references.

**Table S2.** Summary statistics for mitochondrial genome assemblies generated using GetOrganelle.

**Table S3.** Species identification based on comparisons with the Barcode of Life Data System (BOLD).

**Table S4.** Top BLAST matches for specimens with ambiguous genus-level assignments.

## Notes

### Competing Interest Statement

The authors have declared no competing interest.

